# The single amino acid change of R516K enables efficient generation of vesicular stomatitis virus-based Crimean-Congo hemorrhagic fever reporter virus

**DOI:** 10.1101/2025.01.05.631422

**Authors:** Xue Ma, Yanlong Ma, Ziqiao Wang, Fei Feng, Yutong Han, Xinran Sun, Liushuai Li, Zhichao Gao, Manli Wang, Rong Zhang

**Author notes:** Corresponding authors: Rong Zhang, Ph.D. These authors contributed equally to this work (Xue Ma, Yanlong Ma, Ziqiao Wang, Fei Feng).

## Abstract

Crimean-Congo hemorrhagic fever virus (CCHFV) is a medically important tick-borne virus, causing severe hemorrhagic diseases in humans. There is no approved vaccines and therapeutics for CCHFV infection. The study of CCHFV authentic virus requires biosafety level 3 facilities, hindering the research and development of antivirals. Here we report the generation of a recombinant vesicular stomatitis virus (VSV) bearing both CCHFV glycoprotein precursor (GPC) and EGFP reporter (rVSV-CCHFV-GFP). We also find that the acquisition of an unexpected single R516K mutation in the GPC protein enables the packaging of high-titer pseudotyped particles. The replication-competent rVSV-CCHFV-GFP reporter virus resembles the entry properties of authentic virus and allows for rapid assessment of susceptible cell lines, neutralizing antibodies, and host entry factors such as heparan sulfate in fluorescence-based assays. This study provides valuable strategy for packaging of high-titer CCHFV pseudovirus, and the tool generated here can be served for the identification and evaluation of countermeasures against the cell entry of CCHFV.

## Introduction

Crimean-Congo hemorrhagic fever virus (CCHFV) is a tick-borne member of the family *Nairoviridae* in the order *Bunyavirales*, which causes serious hemorrhagic diseases in humans ^1^. The case fatality rate reported by different countries varies from about 5% to as high as 80% ^2^. According to the latest list of pathogens published by the National Health Committee of China, the study of CCHFV has to be conducted in the containment of biosafety level 3 (BSL-3). Given the limited accessibility to studying authentic CCHFV, safe and convenient model systems that can mimic the virus infection like entry steps and be utilized at low biosafety levels such as BSL-2 is needed.

The CCHFV harbors a tri-segmented negative-stranded RNA genome. The large (L) segment encodes viral RNA-dependent RNA polymerase. The middle (M) segment encodes glycoprotein precursor (GPC), and the small (S) segment encodes nucleoprotein (NP) ^3^. GPC is cleaved and modified to generate two structural proteins Gn and Gc, as well as three non-structural proteins GP38, GP85, and GP160^4^. Gn and Gc are responsible for receptor binding and membrane fusion, and are the primary targets of neutralizing antibodies. In addition, mouse studies have shown that the immune response to GPC is critical for protection ^1,5^.

Vesicular stomatitis virus (VSV) is a single-stranded, negative sense RNA virus that can be manipulated at BSL-2. The simple genomic structure of VSV and the capacity to tolerate large heterologous gene fragment make it ideal for exploring the roles of foreign proteins in the context of infectious viral particles^6,7^. Pseudotyped viruses bearing heterologous envelope proteins can mimic the cellular entry of authentic viruses and perform rapid high-throughput diagnostics in BSL-2 facility^8^. Pseudotyped viruses also play important roles in screening of entry-related antivirals, evaluating the activities of neutralizing antibodies, and vaccine development ^9,10^.

The VSV-based system has been successfully used to generate single-cycle or replication-competent multicycle pseudotyped viruses as alternative tools to study the functions of glycoproteins of highly pathogenic viruses, such as Lassa virus^11^, Marburg virus^12^, Ebola virus ^13,14^, Hantavirus ^15–17^, Nipah^18^, SARS-CoV^19^, MERS-CoV^20^, and SARS-CoV-2^21–24^. Although the packaging of single-cycle VSV pseudotyped with the CCHFV-GPC protein with carboxy-terminal deletion of 53 amino acids (GPC^Δ53^) has been reported^25^, the titer is worth to be improved. Also, the replication-competent multicycle VSV bearing the CCHFV-GPC with carboxy-terminal deletion of 14 amino acids (GPC^Δ14^) and some mutations was generated previously^26^, but the expression of both CCHFV-GPC structural protein and reporter genes has not been reported.

In this study, we generated the replication-competent VSV expressing both CCHFV-GPC protein and EGFP reporter, named rVSV-CCHFV-GFP. Interestingly, the deletion of carboxy-terminal 53 amino acids combined with the acquisition of an unexpected single R516K mutation in the GPC during the virus rescue renders the packaging of high titer of single-cycle pseudotyped particle. By using the replication-competent reporter virus rVSV-CCHFV-GFP, the infectivity of a variety of cell lines was evaluated, and antibody neutralization assay was developed and validated. The cell entry property of CCHFV was also examined. The generation of replication-competent rVSV-CCHFV-GFP provides a valuable and convenient tool for studying the CCHFV entry mechanism and inhibitors. The key finding of R516K mutation also helps to package high titer of CCHFV pseudotyped particles.

## Materials and Methods

### Cells

HEK 293T, HeLa, A549, BHK-21, SW13, MDCK, Vero E6, I1-Hybridoma, and Huh7 cells, all were cultured at 37□°C with 5% CO2 environment in Dulbecco’s modified Eagle’s medium supplemented with 10% fetal bovine serum (FBS), 1% MEM Non-Essential Amino Acids Solution, 100 U/mL Penicillin–Streptomycin, 10□mM HEPES and 1% L-glutamine. HEK 293F cells were cultured in SMM293 TII medium. All cell lines were routinely tested for mycoplasma contamination.

### Plasmid construction

Plasmid pB-VSV-GFP transcribing the positive-strand VSV RNA sequences bearing EGFP reporter gene between genes G and L was designed, synthesized, and constructed. Four helper plasmids, pC-VSV-N, pC-VSV-P, pC-VSV-L, and pC-VSV-G, were generated by PCR amplification of each gene from the pB-VSV-GFP plasmid and cloning into pCAGGS vector. The reporter gene of NanoLuc luciferase was PCR amplified and cloned into pB-VSV-GFP plasmid to replace the coding region of G gene to generate the plasmid pB-VSV^ΔG^-Nluc-GFP. The cDNA encoding the open reading frame of CCHFV IbAr10200 strain GPC (NP_950235) was synthesized. The CCHFV-GPC with a deletion of 53 amino acids at the carboxyl terminus (GPC^Δ53^) was PCR amplified and cloned into pB-VSV-GFP plasmid to replace the coding region of G gene. The resulting plasmid was designated as pB-VSV^ΔG^-CCHFV-GPC^Δ53^-GFP. The full-length GPC (GPC^WT^), GPC with a deletion of 14 amino acids at the carboxyl terminus (GPC^Δ14^), GPC^Δ53^, GPC^Δ53^ with a single mutation of L518V or R516K (GPC^Δ53-L518V^ or GPC^Δ53-R516K^), were PCR amplified and cloned into pCAGGS vector for packaging of single-cycle pseudotyped particles. All the plasmids were verified by Sanger sequencing. All the primers used for plasmid construction are available upon request.

### Rescue of replication-competent rVSV-GFP and rVSV-CCHFV-GFP

The GFP reporter virus of recombinant VSV (rVSV-GFP) was rescued using a Vaccinia virus-free recovery system. Briefly, T7RNP-expressing 293T cells (293T-T7) were generated by transducing with lentivirus packaged with pLV-EF1a-T7RNP-IRES-Puro along with psPAX2 and pMD2G plasmids. 293T-T7 cells were seeded in 10-cm dish at the density of 8 × 10^6^ cells, and transfected with pB-VSV-GFP plasmid and four helper plasmids pC-VSV-N, pC-VSV-P, pC-VSV-L, and pC-VSV-G using the CaCl_2_ solution. At 48 h post-transfection, the supernatant was collected, and stored at -80°C. Similarly, for the rescue of recombinant rVSV-CCHFV-GFP virus, 293T-T7 cells were transfected with pB-VSV^ΔG^-CCHFV-GPC^Δ53^-GFP and helper plasmids expressing the N, P, L, and G. After 48 h, the supernatant was collected, and passaged onto BHK-21 cells. The anti-VSV-G neutralizing antibody was added to passaged cells. After passaging three times on BHK-21, the supernatant was harvested as virus stock for use.

### Packaging of single-cycle VSV pseudotyped with different GPCs

The single-cycle VSV^ΔG^-Nluc-GFP (scVSV^ΔG^-Nluc-GFP) virus was rescued, similar to the procedure performed for the rescue of rVSV-GFP by using the plasmid pB-VSV^ΔG^-Nluc-GFP. The scVSV^ΔG^-Nluc-GFP was propagated in cells transfected with pC-VSV-G. To package the single-cycle pseudoviruses bearing different GPCs, 293T cells were transfected with above plasmids expressing the GPC^WT^, GPC^Δ14^, GPC^Δ53^, GPC^Δ53-L518V^, or GPC^Δ53-R516K^ for 24 h, and then infected with scVSV^ΔG^-Nluc-GFP for 2 h. After 3 washes, cells were maintained in culture medium in the presence of anti-VSV-G neutralizing antibody for another 24 h. The supernatant was collected, centrifuged at 3,500 rpm at 4°C for 15 min to remove cell debris, and stored at -80°C. The resulting VSV-based single-cycle pseudoviruses were designated as scGPC^WT^, scGPC^Δ14^, scGPC^Δ53^, scGPC^Δ53-L518V^, and scGPC^Δ53-R516K^.

### Growth curve by plaque-forming assay

BHK-21 cells were incubated with rVSV-GFP or rVSV-CCHFV-GFP (MOI 0.01 2 h). After washing, cells were maintained in 2% FBS culture media, and the supernatant was harvested at various time intervals up to 72 h post infection. Titration was performed using plaque-formation assay on Vero E6 cells. Cells in 96-well plates were incubated with ten-fold serial dilution of viral samples in DMEM with 2% FBS for 2□h at 37°C, and then overlaid with 1% methylcellulose to allow plaques to form. Cells were fixed with 4% PFA for 1 h, washed with PBS, and stained with 1% crystal violet for 15 min, then washed again. Plaques were counted, and viral titer was calculated and expressed as plaque-forming unit per milliliter (PFU/ml).

### Antibody expression

The sequences encoding the anti-CCHFV-Gc neutralization antibody ADI-36121 were extracted from PDB (7KX4), synthesized, and cloned into the corresponding light chain and heavy chain expression vector. HEK 293F cells were transiently transfected with heavy and light chain expression plasmids. For anti-VSV-G neutralization antibody expression, I1-Hybridoma (ATCC CRL-2700) was cultured for 5 days, the supernatant was collected and antibody was purified using protein A beads (GenScript) and dialyzed in PBS.

### Neutralization assay

Antibodies described above or published previously^27^, were serially five-fold diluted and incubated with equal volume of rVSV-CCHFV-GFP (MOI 0.3) for 1 h at 37 °C. Subsequently, the mixture was added to 96-well plates containing monolayer of A549 cells. Plates were incubated for 16 h at 37°C. Cells were collected and fixed with 4% PFA, washed twice with PBS and resuspended in PBS containing 1% FBS. Virus infection was measured by flow cytometry and the IC_50_ (half-maximal inhibitory concentration) value was calculated using a nonlinear regression analysis with GraphPad Prism software.

### *In vitro* acid-induced cell-cell fusion assay

We established an *in vitro* acid-induced quantitative dual-split-reporter-based cell-cell fusion assay as previous described^28–30^, for which the two components of split-GFP and split-Renilla luciferase were individually cloned into pcDNA3.1 vector, resulting in the plasmids DSP1-7 and DSP8-11. Briefly, the effector cells (293T) in 12-well plate were transfected with 0.5 μg pCAGGS empty vector or different expression plasmids of CCHFV-GPC together with 1 μg of one half of dual-split-reporter expression plasmid (DSP1-7), while target cells (Huh-7) were transfected with 1 μg the other half of dual-split-reporter expression plasmid (DSP8-11) using transIT-LT1 reagent. At 24 h post-transfection, effector and target cells were gently detached and an equal number cells were mixed and co-cultured overnight. Then, co-cultured cells were washed with PBS, incubated for 30 min in DMEM Medium at pH 5.2. Following acidic pH treatment, cells were washed with pre-warmed DMEM medium with 10% FBS and incubated for additional period of 24 h. The cells were fixed, stained with DAPI, and the area of GFP-positive cells was quantified using an Operetta High Content Imaging System (PerkinElmer). Twenty representative fields of view were imaged and processed. For luciferase measurement, the EnduRen (Promega, E6481) was added to the plate. The luciferase activity was measured using an automated multimode microplate reader (TECAN Spark) and the SparkControl software.

### Generation of knockout cells

The sgRNA targeting the *B3GAT3, B4GALT7, or LDLR* gene was cloned into plasmid lentiCRISPR v2 (Addgene #52961), and packaged with plasmids psPAX2 and pMD2G in 12-well plate using transIT-LT1 reagent, and the supernatant was harvested after 48 h. The sgRNA sequences are as follows: sgControl, 5’-ACGGAGGCTAAGCGTCGCAA-3’; sg*B3GAT3*, 5’-CCAGAGCCCATACCTGGCAT; sg*B4GALT7*, 5’-CACTACAAGACCTATGTCGG-3’; sg*LDLR*, 5’-TCAAGCATCGATGTCAACGG-3’. A549 cells in 12-well plate were transduced with 100 ul of lentivirus expressing indicated sgRNAs and selected with puromycin for 7 days. The knockout efficiency was verified by probing with anti-B3GAT3 (Santa Cruz #sc-390475) and anti-B4GALT7 (Proteintech #10535-1-AP) antibodies, or analyzing the surface expression of heparan sulfate using flow cytometry.

### Flow cytometry analysis

Cells were fixed with 4% PFA, and then directly subjected to flow cytometry analysis of GFP-positive cells. For surface staining, live cells were incubated with mouse anti-VSV-G (I1-Hybridoma, ATCC CRL-2700), human anti-CCHFV-Gc (ADI-36121 described above), or mouse anti-heparan sulfate (HS) (USBiological #H1890) primary antibody at 4°C for 30 min. Cells were then stained with goat anti-mouse IgG (H + L) (Thermo #A21235), goat anti-human IgG (H + L) (Thermo #A21445), or goat anti-mouse IgM (Thermo #A21238) secondary antibody conjugated with Alexa Fluor 647 for 30 min at 4°C. After two washing, cells were subjected to flow cytometry with Attune NxT (Thermo), and data was processed with FlowJo v10.0.7.

### Heparinase I/III treatment

A549 cells were treated with 2 units/ml of Heparinase I/III (Sigma-Aldrich) in 10% FBS culture medium for 2 h at 37°C. The reduction of heparan sulfate (HS) on the cell surface was determined by flow cytometry as described above. After treatment, the supernatant was removed, and cells were infected with rVSV-GFP or rVSV-CCHFV-GFP at an MOI of 1. After 1 h at 37°C, cells were washed once. At 12 h post-infection, the percentages of GFP-positive cells were measured by flow cytometry and the values were normalized to untreated control.

### Heparin blocking assay

Heparin was dissolved in PBS. rVSV-GFP and rVSV-CCHFV-GFP at an MOI of 1 were pre-incubated with different concentration of heparin at room temperature for 1 h in a total volume of 50 μl, and then added to A549 cells. After 1 h at 37°C, the virus inoculum was removed, and cells were washed once. At 12 h post-infection, the percentages of GFP-positive cells were measured by flow cytometry and the values were normalized to untreated control.

## Results

### Establishment of replication-competent VSV expressing both CCHFV-GPC and EGFP reporter

Previous studies have shown that deletion of 53 residues at the carboxyl terminus of CCHFV-GPC (GPC^Δ53^) is able to improve the packaging efficiency of single-cycle VSV-based pseudotyped particle ^25^. We thus cloned the cDNA encoding the CCHFV-GPC with such a deletion into plasmid pB-VSV-GFP to replace the G gene (Figure 1A). This plasmid was co-transfected with helper plasmids encoding the N, P, L, and G proteins of VSV (Figure 1B). The presence of increasing intracellular GFP fluorescence indicated the successful rescue of replication-competent multicycle VSV bearing both CCHFV-GPC^Δ53^ protein and EGFP reporter (Figure 1C). The rescued recombinant virus, hereafter referred to as rVSV-CCHFV-GFP. In the meantime, as control, we also rescued the recombinant VSV expressing only the EGFP reporter (rVSV-GFP) (Figure 1A to C).

**Figure 1.**
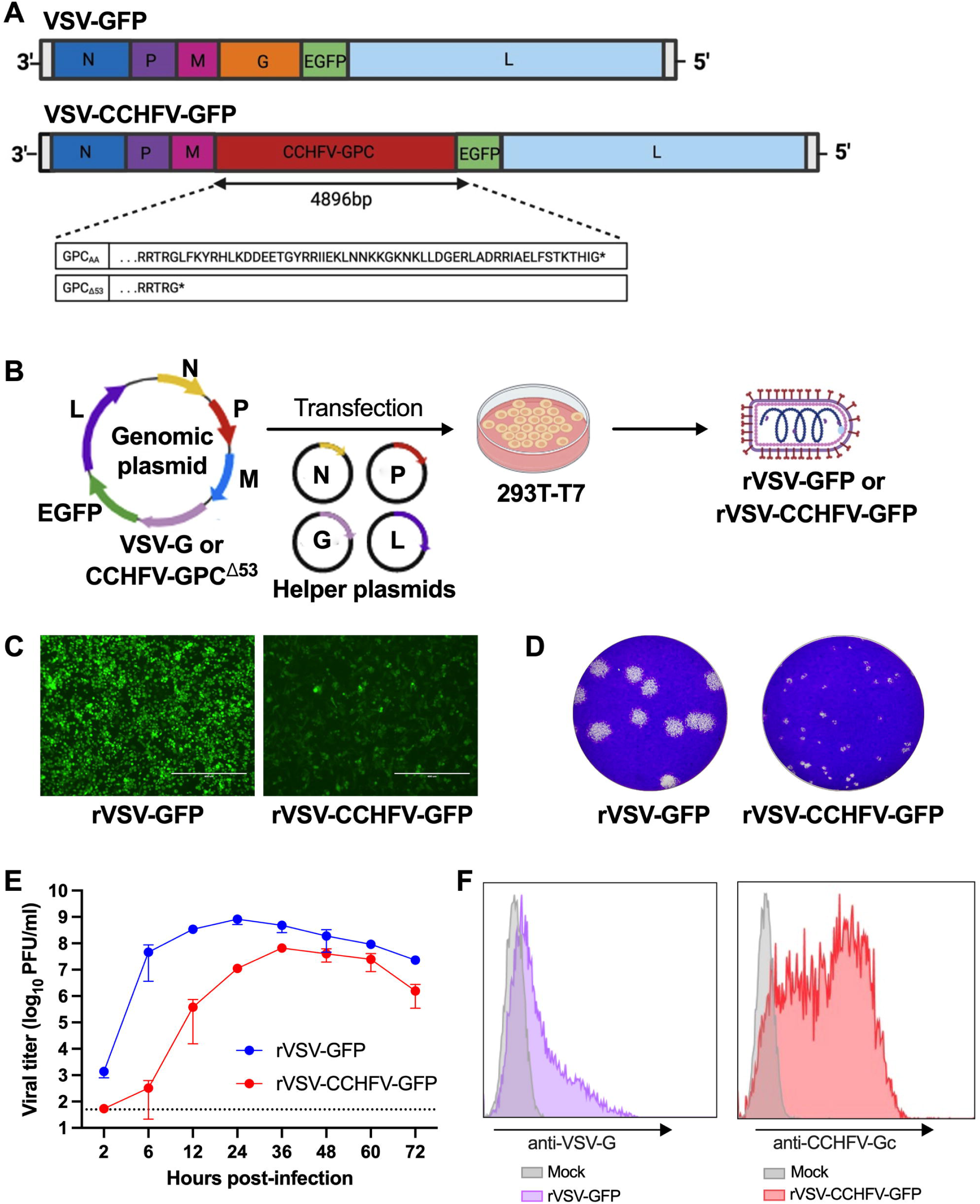
Establishment of replication-competent pseudotyped VSV carrying both CCHFV-GPC and EGFP reporter. (**A**) Schematic of genomic organization of chimeric VSV vectors. EGFP gene was inserted between genes G and L to produce rVSV-GFP reporter virus. The G gene was replaced by CCHFV-GPC with a carboxy-terminal deletion of 53 amino acids (GPC^Δ53^) to generate rVSV-CCHFV-GFP expressing both EGFP and GPC. (**B**) The procedure of rescuing rVSV-GFP and rVSV-CCHFV-GFP viruses. 293T-T7 cells were co-transfected with genomic plasmid pB-VSV-GFP or pB-VSV^ΔG^-CCHFV-GPC^Δ53^-GFP and four helper plasmids expressing the N, P, L, and G. The virus rescue was monitored by detection of GFP signal. (**C**) Representative fluorescent images of BHK-21 cells infected with the rVSV-GFP (MOI 0.01, 24 h) and rVSV-CCHFV-GFP (MOI 0.01, 24 h). Bars, 400 μm. (**D**) Plaque morphology of rVSV-GFP and rVSV-CCHFV-GFP in Vero E6 cells. The plaques at 48 h post-infection were visualized. (**E**) Growth curves of rVSV-GFP and rVSV-CCHFV-GFP in BHK-21 cells (MOI of 0.01). Viral titers were determined by plaque-forming assay. (**F**) The surface expression of VSV-G and CCHFV-GPC protein in mock or virus-infected BHK-21 cells was detected by flow cytometry.

We compared the plaque morphology between rVSV-GFP and rVSV-CCHFV-GFP in Vero E6 cells. The replacement of G protein by CCHFV-GPC^Δ53^ in VSV-GFP virus formed smaller plaques (Figure 1D), an indicative of weaker cytopathic effect of rVSV-CCHFV-GFP than that of rVSV-GFP. The growth kinetics of rVSV-GFP was determined in BHK-21 cells. The maximum titer reached 8.2×10^8^ PFU/ml at 24 h post-infection and declined at later timepoints (Figure 1E). As to the growth kinetics of rVSV-CCHFV-GFP, it grew slower than the rVSV-GFP, and the peak titer is about 6.7×10^7^ PFU/ml at 36 h post-infection in BHK-21 cells (Figure 1E). The surface expression of VSV-G or CCHFV-GPC^Δ53^ protein was verified by flow cytometry (Figure 1F).

### Acquisition of a single R516K mutation in CCHFV-GPC allows packaging of high titer of pseudotyped particle

To identify the potential adaptive mutations occurred in the rVSV-CCHFV-GFP virus rescued above, the CCHFV-GPC coding region was sequenced. Interestingly, it showed that the nucleotide at 1547 of GPC was mutated from G to A, resulting in an amino acid change at position 516 from R to K (R516K) (Figure 2A), which is located at the SKI-1 protease cleavage site (^516^RRLL^519^) between GP38 and Gn in the preGn^31^ (Figure 2B).

**Figure 2.**
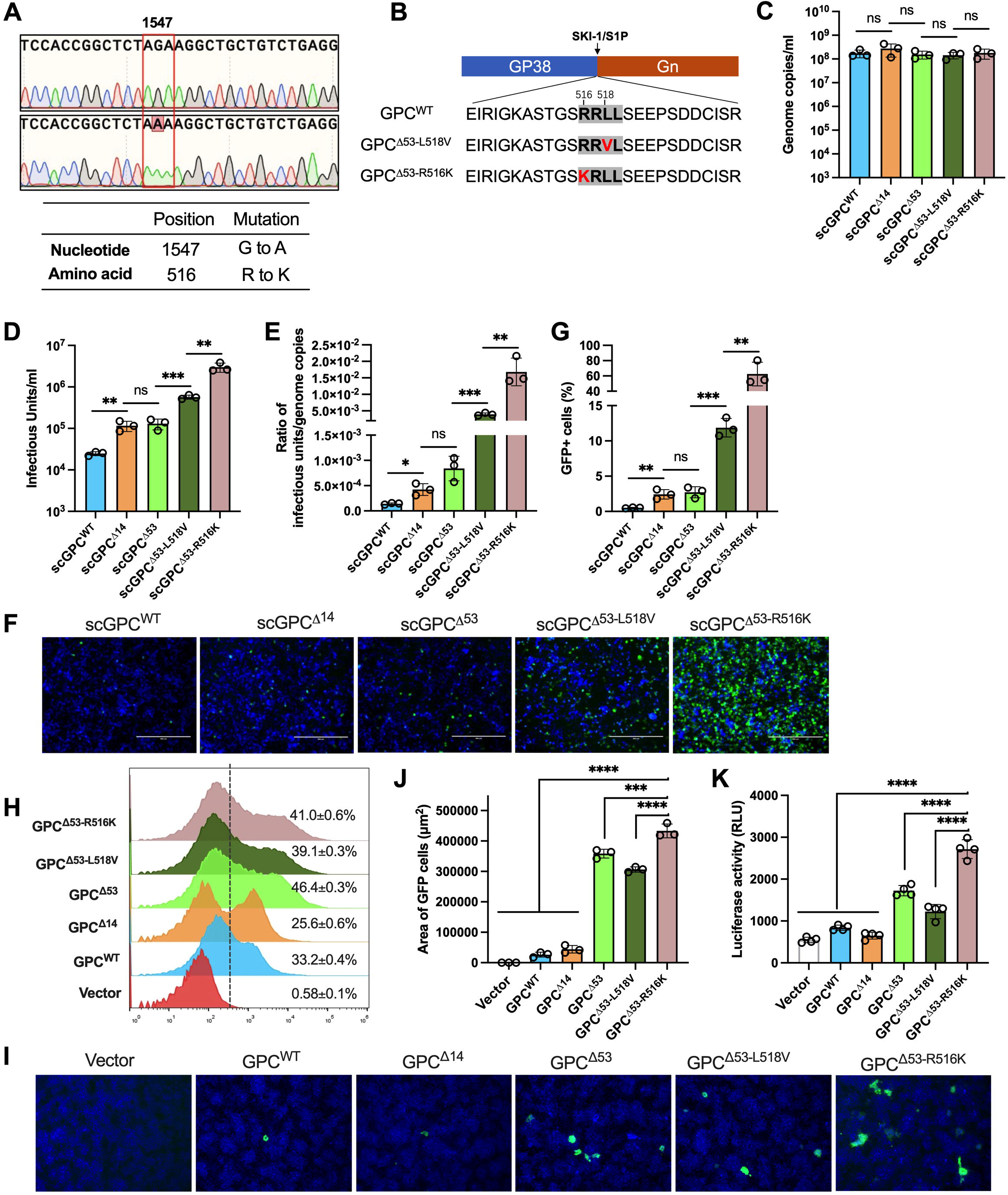
Acquisition of a single R516K mutation in CCHFV-GPC allows packaging of high titer of pseudotyped particle. (**A**) Sanger sequencing of GPC in rescued rVSV-CCHFV-GFP virus. The single nucleotide mutation at position of 1547 and its corresponding amino acid change at 516 from R to K were shown. (**B**) Alignment of the amino acids near the SKI-1/S1P cleavage site “^516^RRLL^519^”. The L518V and R516K mutations are indicated. (**C**) The qRT-PCR was used to quantify the genome copies of single-cycle (sc) pseudotyped particles packaged with different GPCs. (**D**) The flow cytometry was used to quantify the infectious unites of single-cycle pseudotyped particles packaged with different GPCs. The huh7 cells were infected with serial dilution of pseudotyped particles, and the percentage of GFP positive cells was analyzed. The infectious units were calculated according to the number of cells at the time of infection. (**E**) The ratio of infectious units/genome copies was indicated based on the results from figure 2B and C. (**F-G**) The infection efficiency of single-cycle pseudotyped particles packaged with different GPCs in huh7 cells. The same volume of pseudotyped particles was used to inoculate the cells for evaluation. Representative fluorescent images under microscope were indicated (**F**), and the percentage of GFP positive cells was quantified by flow cytometry (**G**). (**H**) The surface expression of Gc protein in 293T cells analyzed by flow cytometry. Cells were transfected with plasmids expressing the full-length, truncated, or mutated GPCs, followed by staining with anti-CCHFV-Gc antibody (ADI-36121). (**I-K**) *In vitro* acid-induced cell-cell fusion assay using the DSP1-7 and DSP8-11 fragments for reconstitution of intact GFP or luciferase. The fused GFP-positive cells were imaged (I) and the area was quantified (J). The activity of reconstituted luciferase after cell fusion was measured (K). C, D, E, G, J, K, unpaired t test; n=3 or 4; mean ± s.d.; *P < 0.05; **P < 0.01; ***, P < 0.001; ****, P < 0.0001; ns, not significant. Data shown are average of three or four independent experiments.

To fully uncover the impact of this R516K mutation and other previously reported deletions or mutations on the packaging of pseudovirus, we constructed a series of transient expression plasmids expressing the full-length wild type GPC without any deletion or mutation (GPC^WT^), GPC with a deletion of 14 or 53 amino acids at the carboxyl terminus (GPC^Δ14^ or GPC^Δ53^), GPCΔ53 with a single mutation of L518V or R516K (GPC^Δ53-L518V^ or GPC^Δ53-R516K^). The GPC^Δ53-R516K^ is identical to the sequences within the recombinant virus rVSV-CCHFV-GFP rescued above. The GPC^Δ14^ with L518V was reported previously in the rescued replication-competent VSV^26^.

The single-cycle VSV-based pseudotyped particles were packaged by transfection of these GPC plasmids into cells, followed by infection with scVSV^ΔG^-Nluc-GFP virus for which the G gene was replaced by Nanoluc luciferase and EGFP reporters. The genome copies of these single-cycle pseudotyped viruses were determined by qRT-PCR, and showed comparable levels (Figure 2C). The infectious units (IU) were quantified in infected cells (Figure 2D). We found that the single-cycle VSV pseudotyped with GPC^Δ53-R516K^ (scGPC^Δ53-R516K^) reaches 3×10^6^ IU, followed by scGPC^Δ53-L518V^ that is 5.7×10^5^ IU. The scGPC^Δ14^ and scGPC^Δ53^ had similar infectious units of around 1.2×10^5^ IU, whereas the scGPC^WT^ was about 2.5×10^4^ IU. The ratio of infectious units/genome copies indicated that more infectious particles of scGPC^Δ53-R516K^ were generated (Figure 2E).

Moreover, the same volume of packaged single-cycle pseudotyped particles was used to infect Huh7 cells for imaging of GFP-positive cells (Figure 2H and G). The average infection efficiency of scGPC^Δ53-R516K^ in Huh7 was 62.5%, whereas the scGPC^Δ53-L518V^ was about 11.9%. Notably, the scGPC^Δ14^ and scGPC^Δ53^ had only around 2.5% of cells infected, and the scGPC^WT^ had about 0.5%. These results demonstrated that 14- or 53-aa deletion at the carboxy terminus of CCHFV-GPC can promote the packaging of VSV virion titer, but the combination of 53-aa deletion with an additional single mutation of R516K dramatically increases the titer, around 5-fold higher than the L518V mutation.

### The amino acid change of R516K increases the fusogenicity of CCHFV-GPC in cells

The surface expression of these CCHFV-GPC proteins with deletions or mutations was assessed, which is the prerequisite for pseudotyped virion budding of VSV. It showed that all the GPCs with different modifications can be expressed on the plasma membrane with percentages ranging from 25% to 46% (Figure 2H). The deletion of 53 amino acids at the carboxy terminus of GPC could obviously increase the cell surface expression, but the mutation of L518V or R516K did not further enhance the expression.

Since the R516K mutation is located in the preGn at the SKI-1 protease cleavage site^31^, we attempted to determine whether the cleavage efficiency of preGn is affected. Unfortunately, the anti-Gn antibodies available were not suitable for Western blotting to probe the cleavage. Likewise, we were unable to investigate if the R516K mutation increases the incorporation of Gn protein into VSV-based pseudovirus particles packaged with the GPC^Δ53-R516K^, as described previously for rVSV-HTNV^32^.

Alternatively, since the GPC structural protein of CCHFV is a class II fusion protein^33^, we then examine if the cleavage of GPC^Δ53^ bearing the R516K could facilitate the cell-cell fusion. We established an *in vitro* acid-induced cell-cell fusion assay using the DSP1-7 and DSP8-11 fragments for which the two components of split-GFP or split-Renilla luciferase were cloned individually^28–30^. The 293T effector cells transfected with both DSP1-7 and different GPCs were co-cultured with Huh7 target cells expressing the DSP8-11, followed by treatment of acid pH of 5.2 to induce the cell-cell fusion. As expected, GPC^Δ53-R516K^ had the highest fusion capacity as indicated by immunofluorescence imaging and quantification of the area of GFP-positive cells (Figure 2I and J). Likewise, measurement of luciferase activity revealed that more cells were fused when GPC^Δ53-R516K^ is expressed (Figure 2K). The increased fusogenicity of GPC^Δ53-R516K^ was possibly due to the efficient cleavage by R516K mutation.

### Evaluation of cell susceptibility and neutralizing antibodies with the rVSV-CCHFV-GFP

We next examined the cell tropism of replication competent rVSV-CCHFV-GFP in different cell lines available in the laboratory. Among the eight cell lines tested, five were highly susceptible to rVSV-CCHFV-GFP infection: BHK-21 (baby hamster kidney), Huh7 (human hepatocarcinoma), Vero E6 (African green monkey kidney), 293T (human embryonic kidney), and A549 (human lung adenocarcinoma). Although the virus replication in SW13 cells (human adrenal adenocarcinoma) was slow at early timepoint as measured by flow cytometry, but the titer in the supernatant could reach about 5.2×10^5^ PFU/ml at 48 h post-infection. In contrast, HeLa (human cervical adenocarcinoma) and MDCK (Madin-Darby Canine Kidney) showed low susceptibility (Figure 3A and B).

**Figure 3.**
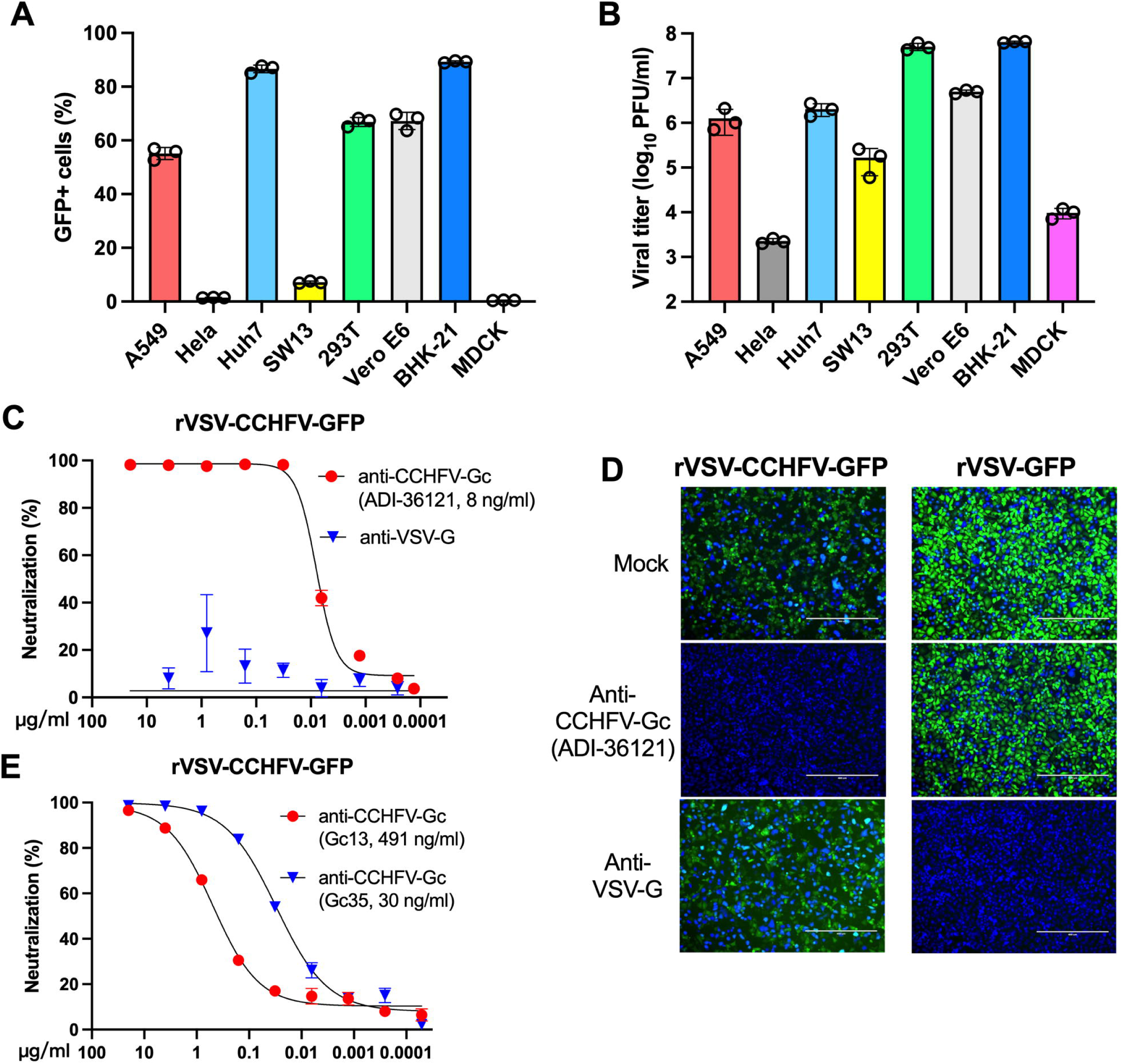
Evaluation of cell susceptibility and neutralizing antibodies with the rVSV-CCHFV-GFP. (**A**) Infectivity of rVSV-CCHFV-GFP on different cell types. Infection efficiency in eight cell lines was measured based on the GFP expression using flow cytometry (MOI 0.3, 20 h). (**B**) Viral titers in culture supernatant of different cell lines were determined by plaque-forming assay (MOI 0.3, 48 h). (**C**) Neutralization of rVSV-CCHFV-GFP virus using anti-CCHFV-Gc (ADI-36121) and anti-VSV-G antibodies (MOI 0.3, 16h) in A549 cells. The percentages of GFP-positive cells were analyzed by flow cytometry. The antibody concentration was shown on the X axis. The IC_50_ value of anti-CCHFV-Gc was indicated in the parenthesis. (**D**) Representative images of A549 cells infected with rVSV-GFP or rVSV-CCHFV-GFP with or without neutralizing antibodies (2 μg/ml) (MOI 0.3, 16h). Bars, 400 μm. (**E**) Neutralization of rVSV-CCHFV-GFP virus using anti-CCHFV-Gc (Gc13, Gc35) antibodies in A549 cells. The percentages of GFP-positive cells were analyzed by flow cytometry. The antibody concentration was shown on the X axis. The IC_50_ values of anti-CCHFV-Gc were indicated in the parenthesis. A-C and E, Data shown are average of three independent experiments and each independent experiment was performed in triplicate.

To evaluate if the replication-competent multicycle rVSV-CCHFV-GFP represents a useful tool for antibody neutralizing assay, we designed a flow cytometry-based fluorescent reporter experiment. The ADI-36121 is a potent anti-CCHFV-Gc human monoclonal neutralizing antibody (NAb) isolated from CCHFV survivors^1,34^. We found that rVSV-CCHFV-GFP can be completely neutralized by ADI-36121, but not by an anti-VSV-G neutralizing antibody (Figure 3C). The IC_50_ values of ADI-36121 against rVSV-CCHFV-GFP was 8 ng/ml (Figure 3C), which is similar to that against CCHFV IbAr10200 tecVLPs reported previously^1^. The blocking effect of NAbs on the infectivity rVSV-CCHFV-GFP or rVSV-GFP virus was indicated by GFP fluorescence as well (Figure 3D). To further evaluate the use of rVSV-CCHFV-GFP for antibody neutralization, we examined two more NAbs reported recently^27^. Both Gc13 and Gc35 antibodies could efficiently neutralize the infection of rVSV-CCHFV-GFP (Figure 3E). The discrepancy of IC_50_ as compared to the literature might be due to the different strains of CCHFV used. Generally, these results suggested that the rVSV-CCHFV-GFP virions resemble the cell tropism and antigenicity of authentic CCHFV.

### Assessment of host entry factors with the rVSV-CCHFV-GFP

Heparan sulfate (HS) is a sulfated glycosaminoglycan (GAG) that is widely expressed on the cell membranes, and can be served as attachment factor for numerous viruses^35^. Both B3GAT3 and B4GALT7 are required for biosynthesis of tetrasaccharide linkage region of proteoglycan and may partially promote the CCFHV infection^36,37^. To verify whether rVSV-CCHFV-GFP can use HS as an attachment factor, we generated *B3GAT3*- and *B4GALT7*-knockout A549 cells. The knockout efficiency of *B3GAT3* and *B4GALT7* was confirmed by Western blotting or analyzing the HS expression (Figure 4A). We found that knockout of *B3GAT3* or *B4GALT7* significantly reduced the infection of rVSV-CCHFV-GFP, but had no or minimal impact on the infectivity of rVSV-GFP (Figure 4B).

**Figure 4.**
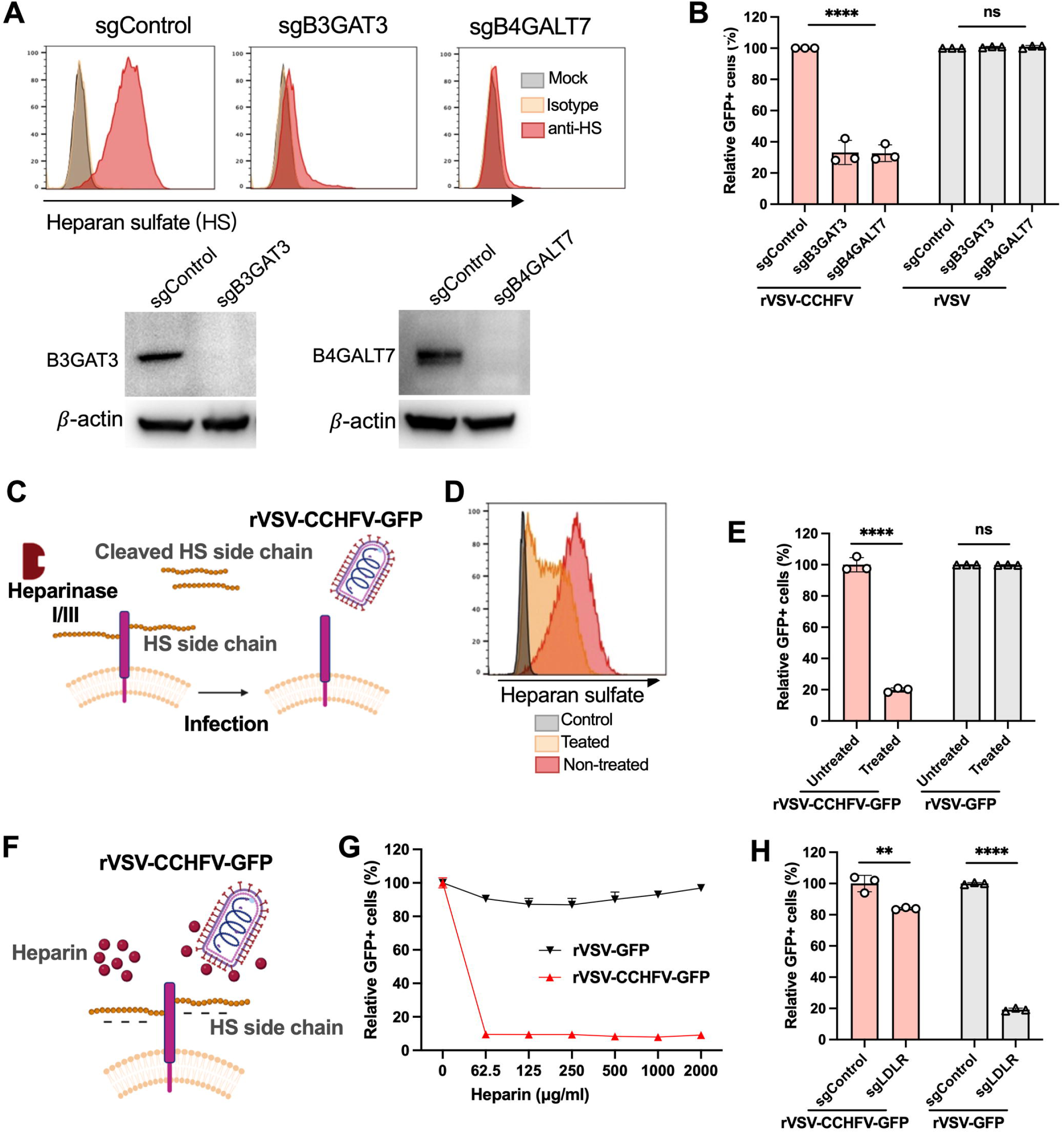
Assessment of host entry factor heparan sulfate with the rVSV-CCHFV-GFP infection. (**A**) The surface expression of heparan sulfate (HS) analyzed by flow cytometry, and editing efficiency of B3GAT3 and B4GALT7 validated by Western blotting. The A549 cells were edited with control-, *B3GAT3*-, or *B4GALT7*-sgRNA followed by staining with anti-HS or isotype antibody, and total cell lysates were harvested and probed with anti-B3GAT3 or B4GALT7 antibodies. (**B**) The A549 cells edited with control-, *B3GAT3*-, or *B4GALT7*-sgRNA were infected with rVSV-GFP or rVSV-CCHFV-GFP , followed by analyzing the GFP-positive cells using flow cytometry. (**C**) Schematic of enzymatic digestion of surface HS side chain in A549 cells by heparinase I/III. A549 cells were pre-incubated with 2 Unit of heparinase I/III for 2 h at 37°C. (**D**) The cleavage efficiency of surface HS by heparinase I/III treatment was analyzed by flow cytometry. (**E**) The infection efficiency of VSV-GFP or VSV-CCHFV-GFP in A549 cells treated or untreated with heparinase I/III. After 2 h of treatment, cells were infected with viruses, followed by flow cytometry analysis of GFP-expressing cells. (**F**) Schematic of pre-incubation of heparin with virus particles. The heparin was diluted in PBS to different concentrations, and pre-incubated with rVSV-GFP or rVSV-CCHFV-GFP for 1 h. Then, the heparin-bound virions were added to A549 cells for 12 h. (**G**) The blocking efficiency of different concentrations of heparin on rVSV-GFP or rVSV-CCHFV-GFP infection, as illustrated in (F). The percentages of GFP-expressing cells were determined by flow cytometry. (**H**) The A549 cells edited with control- or *LDLR*-sgRNA were infected with rVSV-GFP or rVSV-CCHFV-GFP, followed by analyzing the GFP-positive cells with flow cytometry. B, E, G, and H, data shown are average of three independent experiments and each independent experiment was performed in triplicate. B, E and H, unpaired t test; n=3; mean ± s.d.; **P < 0.01; ****, P < 0.0001; ns, not significant.

We then treated WT A549 cells with heparinase I/III to remove HS from the cell surface (Figure 4C). The enzymatic digestion of HS by heparinase I/III was confirmed by flow cytometry (Figure 4D). Consistent with the results from knockout cells, cleavage of HS did not affect the infection of rVSV-GFP, but significantly reduced the infection of rVSV-CCHFV-GFP when compared to the untreated control cells (Figure 4E).

We next assessed the effect of soluble heparin on rVSV-CCHFV-GFP infection. rVSV-GFP or rVSV-CCHFV-GFP virus was pre-incubated with different concentrations of heparin, followed by infection in A549 cells (Figure 4F). The soluble heparin markedly inhibited the infection of rVSV-CCHFV-GFP but not rVSV-GFP (Figure 4G). Thus, these results demonstrated that HS is an important host factor for rVSV-CCHFV-GFP entry.

In addition to the attachment factor heparan sulfate, low density lipoprotein receptor (LDLR) was recently identified as an entry receptor for CCHFV infection^38–40^. To examine if the pseudotyped rVSV-CCHFV-GFP could recapitulate the entry of authentic virus, we generated LDLR-knockout bulk A549 cells with CRISPR sgRNA. Editing of LDLR moderately reduced the infection of rVSV-CCHFV-GFP as shown in Figure 4H. As positive control, infection of rVSV-GFP was significantly decreased in edited cells, as the LDLR serves as a major receptor for the VSV^41^.

## Discussion

The study of authentic CCHFV has to be performed in high level of biosafety laboratory such as BSL-3^42^ in China. The exploration of alternative safe models to mimic the infection will be of significance to accelerate the research and antiviral development of CCHFV. Although a replication-competent multicycle VSV-based virus bearing only the CCHFV-GPC has been described previously^26^, here we generated a VSV-based pseudovirus carrying both CCHFV-GPC and EGFP that can replicate robustly. The reporter virus with EGFP expression developed here could significantly simplify the detection procedure and be applied to high-throughput screening assays. The virus of rVSV-CCHFV-GFP could infect various cell lines from different species, exhibiting a wide range of cell tropism. The results were generally consistent with the cell tropism assessed with authentic CCHFV^43^. The use of viral strains, timing, detection methods, and infection doses may contribute to the minor difference in susceptibility. We also applied our rVSV-CCHFV-GFP reporter virus to evaluation of antibody neutralization activity. HS has been identified as an important host dependency factor for many viruses, such as RVFV, ZIKV, SINV, and SARS-CoV-2, by facilitating the initial attachment of virions to host cells^37,44–46^. By using rVSV-CCHFV-GFP as model virus, we verified the role of HS in promoting CCHFV infection^37^. Likewise, we validated the new receptor LDLR with rVSV-CCHFV-GFP^38–40^.

In the current study, the truncated GPC that contains a 53-amino acid deletion in the carboxyl terminal region of Gc was selected to rescue the chimeric rVSV-CCHFV-GFP. As previously reported^25^, the truncation of carboxy-terminal CCHFV-GPC may facilitate the incorporation of GPC protein into VSV particles. It is true that, when compared to the full-length GPC^WT^, deletion of 14 or 53 amino acids increased around 5 folds of virus titer as demonstrated by packaging of the single-cycle pseudotyped particles. It appears that deletion of either 14 or 53 amino acids has similar titer. Notably, the introduction of a single mutation of L518V described previously^26^, had about 5-fold increase of titer, but the mutation of R516K identified in this study surprisingly enhanced the titer of single-cycle pseudovirus by over 25 folds as compared to the GPC with a 53-amino acid deletion. These mutations occurred at the SKI-1 protease cleavage site (^516^RRLL^519^) of preGn. As compared to CCHFV GPC, the glycoprotein precursor of Lassa virus also has the same “RRLL” cleavage site for SKI-1 protease, and the mutation of “RRLL” to “RRLA” attenuates the efficient cleavage^47^. Interestingly, the first “R” (P4 position) of “RRLL” is absolutely conserved in all of the New World and Old World arenaviruses, implying the critical function of this residue for recognition and cleavage by SKI-1^48^. Indeed, a study reported recently showed that R516S mutation abrogates the SKI-1 cleavage between GP38 and Gn of CCHFV^49^. Thus, whether the amino acid substitution at position 516 from R to K at the SKI-1 protease cleavage site of preGn facilitates the proteolytic cleavage and maturation of GPC protein thereby facilitating the appropriate assembly of VSV virions or not, remains to be further investigated.

In conclusion, we have successfully generated a replication-competent rVSV-CCHFV-GFP reporter virus, and identified the key single amino acid change of R516K for packaging of pseudovirus particles with high titer. Although the distribution, conformation, and density pattern of GPC protein with the carboxy-terminal deletion of 53 amino acids and a mutation of R516K incorporated into rVSV-CCHFV-GFP virion may not fully reflect its “natural” state on the surface of authentic virus particles^50^, and the VSV-based pseudovirus can only mimic the entry stage of infection lifecycle, it provides a safe and valuable alternative tool to study the roles of CCHFV-GPC in mediating entry into host cells, to screen and assess neutralizing antibodies, and to develop vaccines and entry inhibitors.

## Acknowledgements

This project was supported by the Shanghai Municipal Science and Technology Major Project (ZD2021CY001), Program of Shanghai Academic/Technology Research Leader (22XD1420600), Non-profit Central Research Institute Fund of Chinese Academy of Medical Sciences (2023-PT310-02), and Shenzhen Medical Research Fund (SMRF No. B2302029). Some of the figures were created using BioRender.com.

## Data Availability

The authors declare that all relevant data supporting the findings of this study are available within the paper. Any other data of this study are available upon request.

## Author Contributions

Y.M., X. M., Z.W., F.F., and R.Z. designed and performed the experiments. Y.H., X.S., Z.G. provided technical or material support. Y.M., X. M., F.F., R.Z performed data analysis. X.M. and R. Z. wrote the the manuscript.

